# Robust and cost-efficient single-cell sequencing through combinatorial pooling

**DOI:** 10.1101/2024.11.22.624460

**Authors:** Johannes Gawron, Luís Cunha, Nico Borgsmüller, Niko Beerenwinkel

## Abstract

Single-cell sequencing is widely used to study molecular cell-to-cell heterogeneity. Even though the cost of sequencing has dropped throughout the last decades, single-cell assays remain expensive, because they require strategies to index molecules by cells. The high costs of indexing can be mitigated by pooling samples prior to sequencing library preparation. Computational methods have been developed to leverage molecular features that are distinct between different samples to separate the pools into distinct datasets. However, since all multiplexed samples are processed in the same way, information on the origin of each demultiplexed dataset is lost. To map datasets to their sample of origin, additional information such as molecular indexing or additional genotyping is needed. Here, we propose a class of experimental designs that allows identifying the sample of origin of each demultiplexed dataset, only relying on the genetic profiles of the samples and the composition of pools. Our approach is based on splitting and pooling samples in specific combinations. We find a most cost-efficient experimental design in this class and prove its optimality. We present a dynamic programming algorithm to iteratively simplify an optimal experimental design by breaking it into several independent designs while maintaining optimality. Furthermore, we propose a subclass of experimental designs which allow robust sample identification even under partial failure of the experiment and present a provably optimal design in this subclass. We provide an implementation for automatic sample identification under these optimal combinatorial pooling strategies and demonstrate its functionality in a simulation study.

## 1 Introduction

Sequencing the genome or transcriptome on the level of a single cell has become a wide-spread approach to study heterogeneity among cells. For example, in cancer research, single-cell sequencing has enlightened the role of co-evolution and treatment resistance of genetically distinct tumour clones under selection [21,32,36], enabled the detection of prognostic biomarkers [29,31], and is considered as a diagnostic tool for everyday clinical decision making in cancer treatment [11].

However, the quality of single-cell datasets suffers from noise and contamination with doublets, the accidental and undetected sequencing of two cells as one cell [12], with typical doublet rates below 8% [7,20]. High costs of single-cell genomics and transcriptomics assays constitute another major limitation of these technologies and hampers broader applications in research and the clinics [15,23,26].

Costs can be reduced by mixing the tissue material to obtain a pool of different samples prior to library preparation and sharing its costs among the samples. This procedure is called sample multiplexing [13,30]. After sequencing, one obtains a single-cell dataset, where the cells originate from several donors.

This raises two computational challenges: First, cells from a multiplexed dataset need to be clustered by samples of origin and doublets need to be distinguished from true single cells to be able to analyse datasets by sample, a procedure called demultiplexing [12]. Second, each of the resulting clusters need to be correctly annotated by its sample of origin. We refer to this task of finding a mapping between demultiplexed datasets and their sample of origin the sample identification problem. There are two different strategies that tackle both problems.

The first approach assumes that all samples in a pool come from a distinct donor of origin. It harnesses the abundance of genetic variability of single-nucleotide polymorphisms (SNPs) to split the dataset, both in the setting of scRNA-seq data [35,12,13,9,10,6], and scDNA-seq data [2,19]. Since all samples undergo the same library preparation and sequencing procedure, it is a priori unclear which datasets corresponds to which of the donors (Figure 1**a**). The sample identification problem is usually addressed by additional genotyping using, for example, a SNP array [16,14], on additional tissue material from the donors. The identified SNPs in each demultiplexed dataset are then used as features for the clustering and matching.

**Fig. 1.**
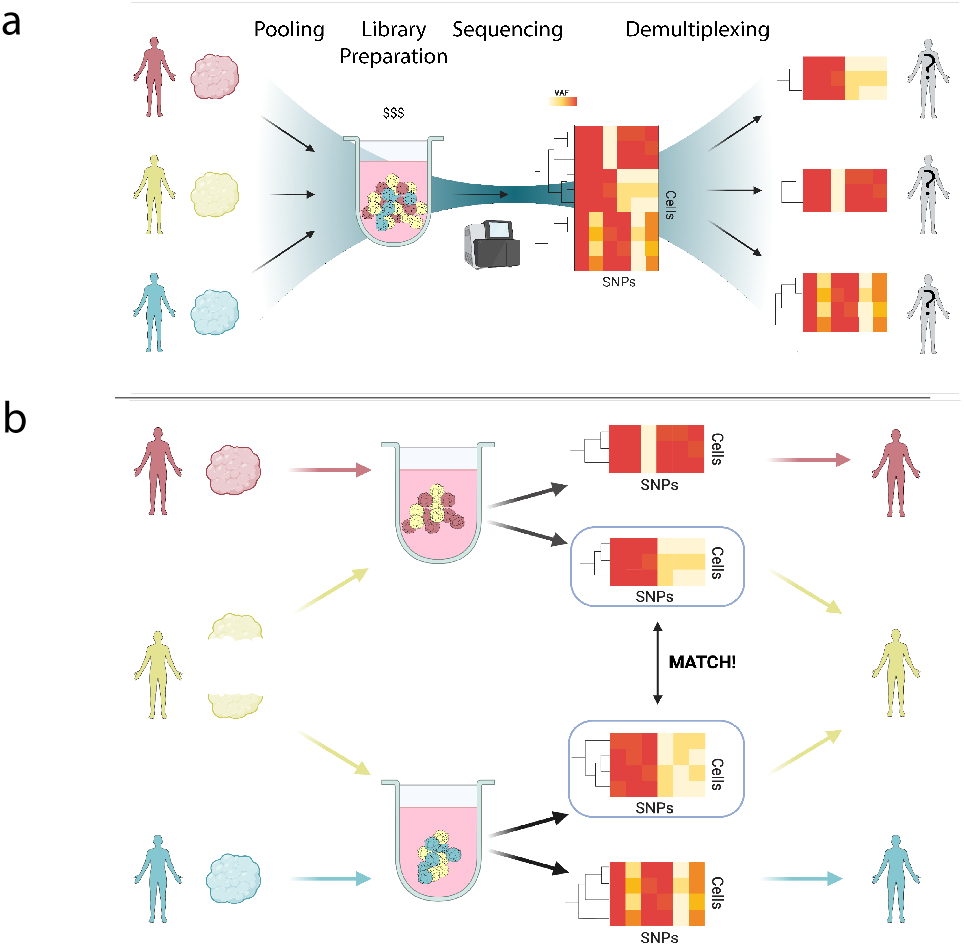
To reduce costs of single-cell sequencing, sample multiplexing is performed and a pool of cells from different donors is sequenced jointly. Demultiplexing algorithms allow to distinguish the different samples from each other. **a**. However, since all samples are processed in the same way, the identity of each sample’s donor of origin is lost. **b**. To overcome this problem, each sample is split and pooled in a unique pattern. By matching genotypes of demultiplexed samples, the pooling pattern can be recovered and the donor of origin identified.

The second approach experimentally introduces sample specific features prior to pooling by labelling cells with sample-specific oligonucleotide unique molecular identifiers (UMIs) [28,3,18,5,27,8]. The demultiplexing algorithms work by estimating UMI count distributions for cell populations that are considered UMI-negative or ambient RNA, and compares the UMI count of each cell to this distribution [1,34,4,17]. Barcoding-based procedures allow to map the demultiplexed datasets directly to the donor of origin using the UMI labels, but require the use of additional reagents and preprocessing prior to pooling.

In this work, we propose a method for sample identification after demultiplexing that does not rely on any additional genotyping or barcoding, when the samples are taken from unrelated donors. The method is based on physically splitting each sample into several subsamples prior to multiplexing them into a different pool according to a predefined rule. This rule, the pooling strategy, requires that no two donors are mixed into the same subset of pools. Each pool is sequenced and the resulting dataset is demultiplexed based on the variability of SNPs. We infer a consensus genotype on each sample and compare the pairwise genetic distance of demultiplexed samples from different pools to find genetically identical subsamples. We conclude that these must originate from the same donor, since unrelated donors have different genotypes. Since at most one donor is associated with each subset of pools, the donor’s identity can be uniquely determined by this piece of information. Hence, comparing genotype profiles and correctly determining the subset of all pools associated with a genotype profile is sufficient to determine the donor of origin (see Figure 1**b**).

Errors may occur through accidentally incorrect execution of the pooling strategy or imperfect demultiplexing, leading to cross-contaminated genotypes. To account for this possibility, we propose a method that allows sample identification under partial unavailability of data (for example, due to partial failure of the experiment). This approach is robust to incorrect demultiplexing of one pool or ambiguous sample matching. There is a trade-off between achieving higher cost saving and more robust sample identification, hence we explore the best possible cost savings under the above described procedures and constraints. Using this theory, we implemented a method that generates a formal description of an optimal pooling strategy under given experimental conditions. As a second step, it takes demultiplexed genotype profiles for each dataset in each pool and outputs their sample of origin, solving the sample identification problem.

## 2 Results

In Subsection 2.1, we provide an overview of the method, introduce mathematical formalism, and state the main theoretical results. In Subsection 2.2, we demonstrate the method on a dataset of in-silico multiplexed samples.

### 2.1 Theoretical Results

We define mathematical objects that describe the procedure of sample multiplexing and encode the necessary properties to accomplish sample identification. We present an objective function that quantifies the cost savings of a pooling strategy and analytically find an optimum solution of this function. Finally, we explore some related formulations of the optimisation problem under additional robustness and cohort size constraints.

#### Multiplexing schemes

Multiplexing involves physically mixing cells from different samples. Each sample is associated with a pool, meaning that the pool contains the sample. Samples can be split and mixed into several pools, in which case we associate a sample with a subset of pools. This process can be described by a map between two sets, the set of samples and the set of subsets of pools. To ensure that all of the samples are sequenced, every sample must be associated to at least one pool. To be able to distinguish samples by subsets of pools, we need to ensure that no two distinct samples are pools in the same subset of pools. We summarise these assumptions in the following definition.

##### Definition 1.

*Let S, Ω be finite sets. We call a map Φ* : *S → 𝒫(Ω*) *from S into the powerset of Ω a* multiplexing scheme *if it is injective (i*.*e*., *Φ*(*s*) = *Φ*(*s*^*′*^) *implies s* = *s*^*′*^*) and none of the element of S map to the empty set. We call S the set of* samples *and Ω the set of* pools.

##### Example 1.

In the situation of Figure 2**a**, three samples are all associated with one pool. Representing the samples as *S* = {*s*_1_, *s*_2_, *s*_3_} and the pool by *Ω* = {*ω*}, the procedure translates to the map *Φ* : *S → 𝒫(Ω*) defined by *Φ*(*s*_*i*_) = {*ω*} for each *i*. However, all three samples map to the same subset of pools. Hence, *Φ* is not a multiplexing scheme.

**Fig. 2.**
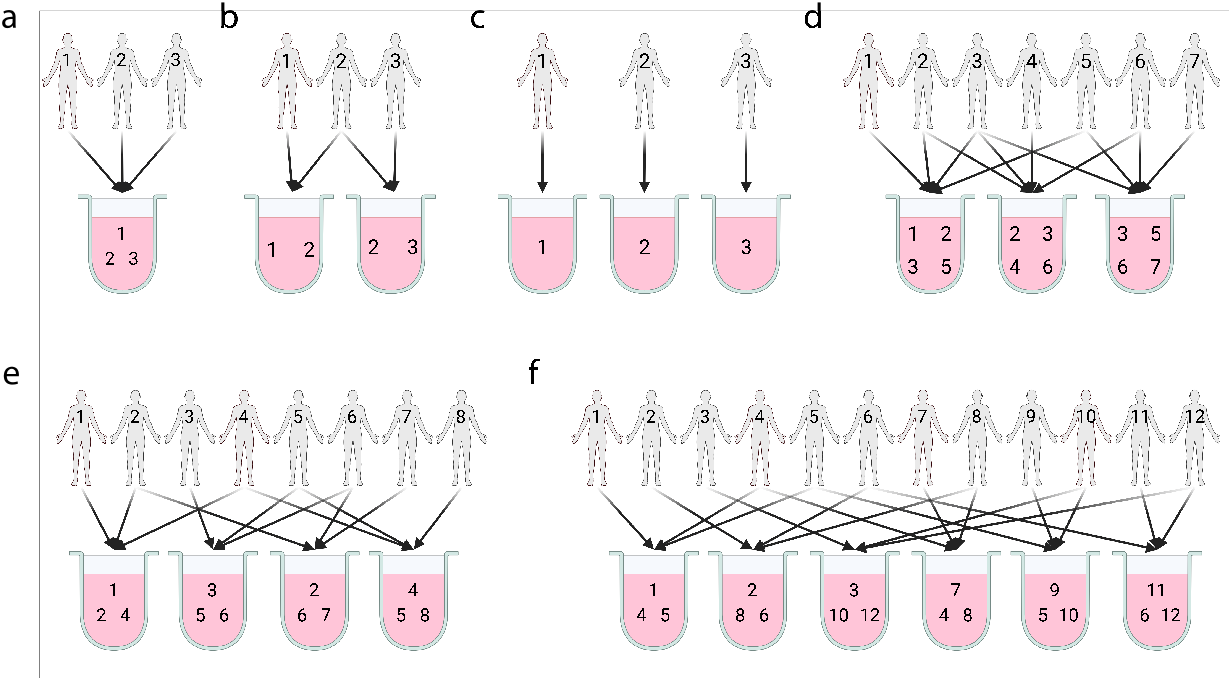
Examples of sample multiplexing. **a**. Three samples are pooled into one pool. Sample identification based on the structure of the pooling is not possible. **b**. Three samples are pooled into two pools. There can be only one sample that appears in both pools (sample 2), only the first pool (sample 1) and only in the second pool (sample 3). **c**. Not mixing samples at all allows sample identification directly, but without any benefits of pooling. **d**. A total of seven samples can be pooled into three pools such that the pooling strategy alone allows sample identification. It requires that four samples are pooled at once. **e**. Eight samples can be pooled into four pools at a maximal pool size of three. **f**. Similarly, twelve samples can be pooled into six pools at a maximal pool size of three. It cannot be decomposed into two independent multiplexing schemes.

##### Example 2.

Consider Figure 2**b**. As opposed to the previous example, two distinct pools are used. We write *Ω* = {*ω*_1_, *ω*_2_}. The first sample *s*_1_ is mapped to the first pool only, *Φ*(*s*_1_) = {*ω*_1_}, the second sample is split and mixed into both pools, *Φ*(*s*_2_) = {*ω*_1_, *ω*_2_}, and the third sample is mapped to the second pool only, *Φ*(*s*_3_) = *ω*_2_. Since each of the subsets of pools is associated with not more than one sample, the map is injective. So *Φ* defines a multiplexing scheme.

The performance of demultiplexing algorithms drops with increasing sample number [37]. However, the number of samples multiplexed in one pool is limited in practice, and usually less than ten [28,9,8]. To account for this restriction, we will study only those multiplexing schemes that lead to pool sizes of a given maximal size. The set of samples that are pooled into a specific pool *ω ∈ Ω* by a multiplexing scheme *Φ* : *S → 𝒫(Ω*) is the fibre *Φ*^*−*1^(*ω*) = {*s ∈ S* | *ω ∈ Φ*(*s*)}. To define pooling strategies with limited pool size, we ask for the fibres of *Φ* to be of bounded cardinality:

##### Definition 2.

*Let Φ* : *S → 𝒫(Ω*) *be a multiplexing scheme. We say Φ is a k*-scheme *if, for each ω ∈ Ω, the set Φ*^*−*1^(*ω*) *has at most k elements*.

By this definition, 1-schemes correspond to library preparation without sample multiplexing (Figure 2**c**), Example 2 defines a 2-scheme, and Figure 2**d** defines a 4-scheme with 7 samples and 3 pools.

**Efficiency**. If up to four samples are pooled together, both Figure 2**b** and **d** solve the sample identification problem, but at different costs per sample. We describe this formally in terms of multiplexing schemes.

##### Definition 3.

*1. The* efficiency *of a multiplexing scheme Φ*: *S → 𝒫(Ω*) *is*

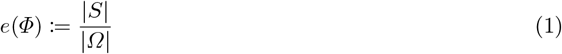

2. For *k* ∈ ℕ_≥1_, the maximum efficiency of any k-scheme is

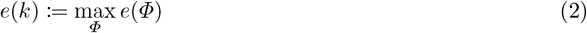

*where the maximum is taken over all k-schemes*.

We ask for the optimal way to multiplex samples so that sample identification is possible and never more than *k* samples are pooled together.

##### Theorem 1.

*The highest achievable efficiency for a k-scheme is*

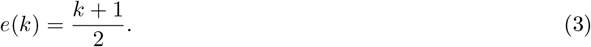

*Furthermore, there is an optimal k-scheme with k pools and* 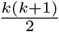 *samples, and it can be constructed explicitly*.

*Sketch of Proof*. The problem is solved analytically by placing *k* arbitrary samples into a single pool each, and splitting the remaining 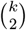 samples into two and assigning each to a different pair of pools (see Figure 3). The complete proof can be found in the supplementary information. □

**Fig. 3.**
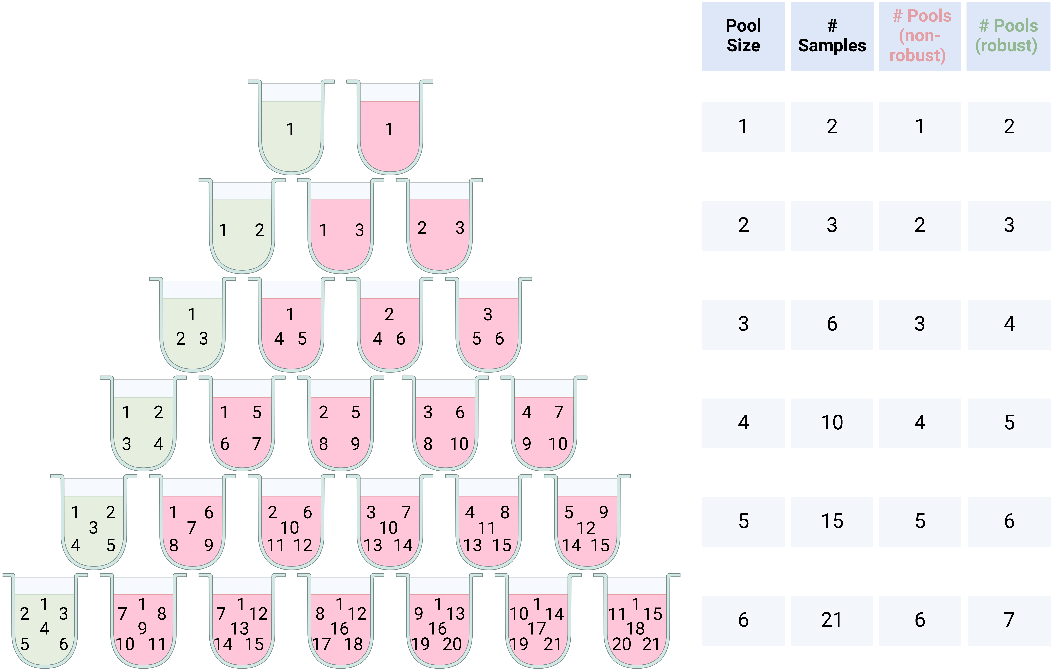
Examples for optimal *k*-schemes (Theorem 1) for *k* = 1, …, 6 (in pink). Samples are indicated by integers and pools represented graphically. A mapping from a sample to a pool is represented by placing the sample number into the pool. Each optimal *k*-scheme can be turned into an optimal robust *k*-scheme (Theorem 3) by adding one more pool (shown in green).

Figure 2**b** is an example of an optimal 2-scheme. Since constructing an optimal *k*-scheme involves enumerating all 1-element and 2-element subsets of a *k*-element subsets, the computational complexity of this construction is *O*(*k*^2^), and *k* is typically small (*k* ≲ 10).

#### Optimal multiplexing schemes for a fixed sample size

Optimal *k*-scheme experiments can be applied independently to batches of 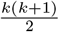 samples, so Theorem 1 is applicable whenever the cohort consists of a multiple of 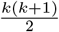 samples. When a cohort of a different size is to be sequenced, optimal efficiency may not be achieved. In the following, we quantify the best achievable efficiency under additional sample size constraints. With fixed sample size, optimizing the efficiency is equivalent to minimizing the number of pools.

##### Theorem 2.

*Let k ∈* ℕ_*≥*1_, *n ∈* ℕ_*≥*1_ *and* 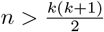.

1. *Let k be odd. Then the smallest number of pools to sequence n samples in a k-scheme is given by*

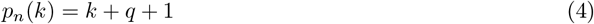

*where q is the (unique) quotient to the Euclidean division equation with remainder r:*

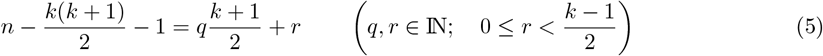
2. *Let k be even. Then the smallest number of pools to sequence n samples is given by*

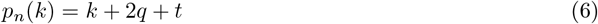

*where q is the (unique) quotient of the Euclidean division equation with remainder r:*

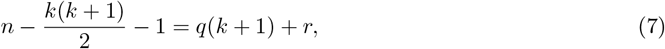

*where, if* 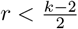 *then t* = 1, *otherwise t* = 2.

*Consequently, the highest achievable efficiency is*

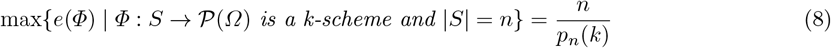

*Sketch of Proof*. The proof proceeds by induction over *q*. Starting with the result of Theorem 1, we show that each added pool allows to multiplex 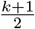 samples if *k* is odd, and 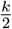 samples if *k* is even and *q* is even, and 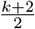 samples if *k* is even and *q* is odd. The proof can be found in the supplementary information. □

In the above theorem, we specify the smallest possible number of pools for a fixed number of samples to allow sample identification with a pool size limited to *k*. Solving the division equation for *n* specifies the maximal number of samples for a *k*-scheme with a given number of pools.

The statement of Theorem 2 assumes that the cohort to be sequenced consists of at least 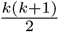 samples. If the cohort is smaller, we find *k*-schemes with a brute force algorithm: Since *k* is small (*k <* 10), we can simply search all possible maps from the set of samples to non-empty subsets of pools and check that the map is a valid *k*-scheme. The smallest number of pools for which this is the case is a solution to the optimisation problem.

##### Example 3.

Figure 2**d** provides an example of an optimal 4-scheme with 7 samples. It requires 3 pools. Figure 2**e** shows an optimal 3-scheme for 8 samples. According to Theorem 2, the minimal number of pools is computed by solving the Euclidean division equation 8 *−* 6 *−* 1 = 4*q* + *r* (*q* = 0, *r* = 1) and calculating *p*_8_(3) = 3 + 0 + 1 = 4.

#### Robust multiplexing

In practice, to be able to perform sample identification for an optimal *k*-scheme, the data must be available for all pools and there must be no ambiguity when matching samples of the same genotype. For a more error-robust approach, we now constrain multiplexing schemes to allow sample identification if the data of an entire pool is unavailable or lost.

##### Definition 4.

*We call a scheme Φ* : *S → 𝒫(Ω*) robust *if, for any ω ∈ Ω, the map* 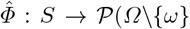) *defined by s 1→ Φ*(*s*)*\{ω} is a multiplexing scheme*.

##### Example 4.

None of the examples in Figure 2 are robust *k*-schemes. In **b**, if the left pool is removed, then the resulting map sends sample 1 to the empty set and this is not a *k*-scheme. In **d**, if the leftmost sample is removed, sample 1 will not be sequenced, and sample 2 is not distinguishable from sample 4 based on the pooling strategy.

Akin to Theorem 1, we have the following efficiency result for robust *k*-schemes.

##### Theorem 3.

*Let k ∈* ℕ_*≥*1_. *The best achievable efficiency for a robust k-scheme is*

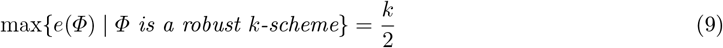

*Furthermore, there is an optimal robust k-scheme with k* + 1 *pools and* 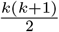 *samples, and it can be constructed explicitly*.

*Sketch of Proof*. The main idea of the proof starts with an optimal *k*-scheme as constructed in the proof of Theorem 1 and adds one additional pool that contains all samples that have not been split. The complete proof can be found in the supplementary information. □

Optimal *k*-schemes, for pool sizes *k* = 1, …, 6, in the robust and non-robust case are displayed in Figure 3.

#### Reducible multiplexing schemes

Using Theorem 2 and the inductive procedure in the proof, optimal *k*-schemes can be constructed for all given sample sizes. In general, it may result in complex procedures where many pools need to be prepared simultaneously. Often, the complexity can be reduced by breaking the scheme into two independent smaller multiplexing schemes. Such a scheme is required to satisfy that a given subset of samples can be fully pooled and sequenced using only a subset of the pools. In other words, the sequencing experiment is broken down into two independent experiments.

##### Definition 5.

*Let Φ* : *S → 𝒫(Ω*) *be a k-scheme. The we call Φ* reducible, *if Ω is the disjoint union of sets Ω*_1_ *and Ω*_2_ *such that, for all s ∈ S*,

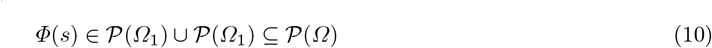

If *Φ* : *S*_1_ *→ 𝒫(Ω*_1_) and *Ψ* : *S*_2_ *→ 𝒫(Ω*_2_) are two *k*-schemes, then the assignment

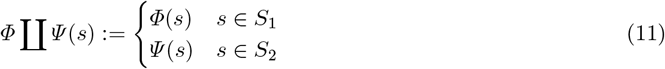

defines a reducible *k*-scheme.

In the following, *p*_*i*_(*k*) is the minimal number of pools to multiplex *i* samples in a *k*-scheme. Starting with an arbitrary *k*-scheme for fixed sample size such as provided in Theorem 2, we can detect reductions of *k*-schemes using a dynamic programming algorithm. The procedure is summarised in Algorithm 1. We define *Φ*_*i*_(*k*) as an optimal *k*-scheme for *i* samples. By iterating through the number of samples, we update the value of *Φ*_*i*_(*k*) if it can be reduced to two *k*-schemes *Φ*_*j*_(*k*) and *Φ*_*i−j*_(*k*). The time complexity of the algorithm is *O*(*n*^2^).

##### Example 5.

The demultiplexing scheme in Figure 2**f** is an irreducible 3-scheme pooling 12 samples into 6 pools. It can be replaced by a 3-scheme that is reducible into two optimal 3-schemes as in Figure 3.

### 2.2 Computational results

We implemented our method to computationally solve the sample identification problem. It takes the number of samples *n* and the maximal pool size *k* as input and generates a multiplexing scheme. In Figure 5, we summarize the cost savings per sample as a function of *n* for different pool sizes *k*. Costs are reported relative to the non-multiplexed case. For example, for 30 samples and a maximal pool size of 3, costs are halved, while for a maximal pool size of 6, costs are reduced by two thirds.

**Fig. 4.**
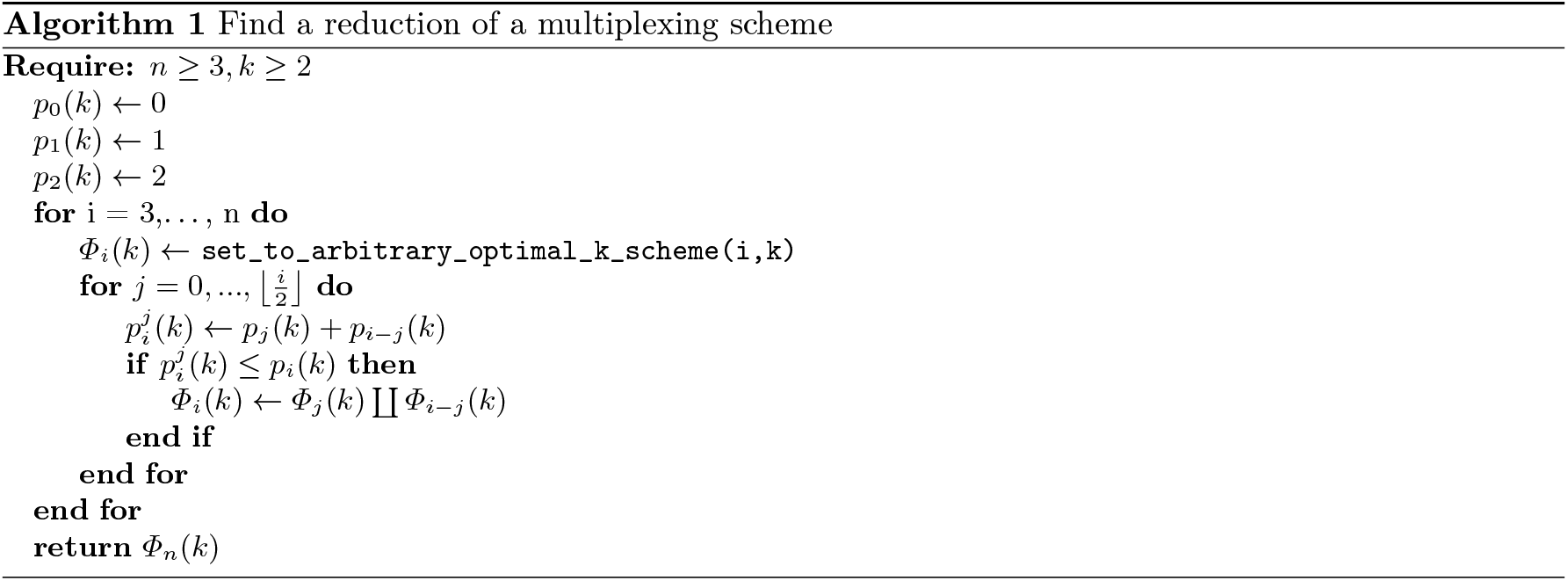
Dynamic programming algorithm to identify reductions of a multiplexing scheme. By iteratively increasing the sample size, the algorithm checks if a multiplexing scheme can be replaced by two independent multiplexing schemes of the same efficiency.

**Fig. 5.**
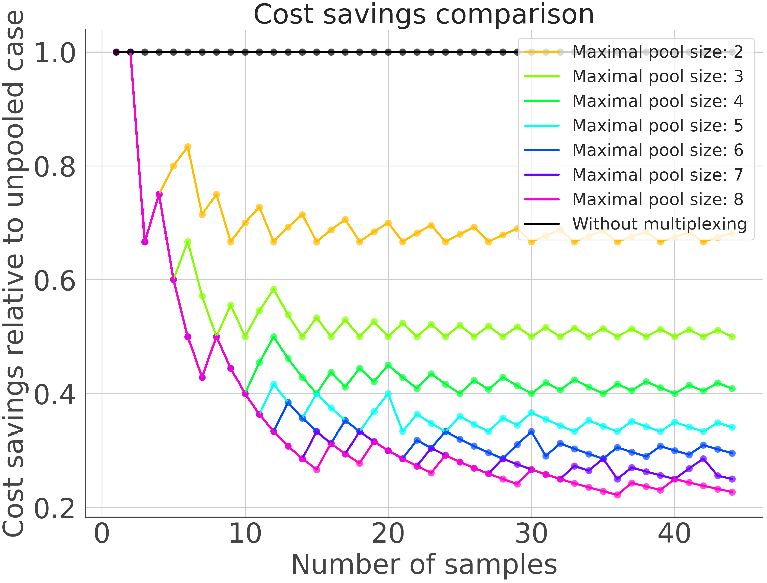
Cost savings relative to the non-multiplexed case. The x-axis represents the cohort size, the y-axis the cost of library preparation per sample normalized by the cost of library preparation under an optimal *k*-scheme. Varying colours indicate different values of *k*. In the limit of large sample numbers, the costs converge to the best achievable efficiency (Theorem 1).

Furthermore, our method takes as input the genetic profiles of the demultiplexed datasets given by a matrix *X*_*P*_ *∈* [0, 1]^*M×N*^ for each pool *P*, where *M* is the number of demultiplexed datasets, *N* is the number of loci, and each entry (*X*_*P*_)_*ij*_ is the variant allele frequency of sample *i* in pool *P* at locus *j*. Its output is a mapping from each dataset in each pool to its sample identity.

To accomplish sample identification, we consider all pairs of pools for which we expect to find a sample that was split into these, and compare the pairwise genetic distance between pairs of datasets from the two different pools (see Subsection 3.1). For each candidate pair of pools, we calculate the ratio of the smallest and the second smallest pairwise distance and consider the signal in this pool to be strong when this ratio is small. We iteratively match the samples with the smallest distance from the pair of pools with the strongest signal (see Figure 6).

**Fig. 6.**
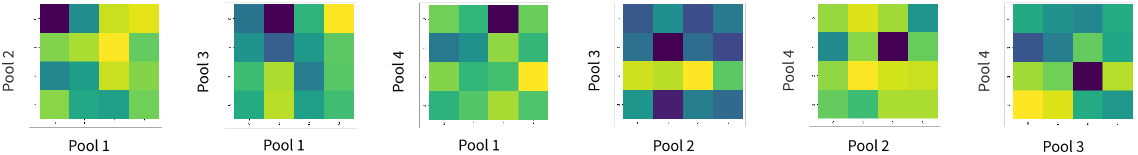
Illustration of computational methods. Sample matching is accomplished by comparing the pairwise genotype distance in demultiplexed datasets (purple indicates small distance, yellow indicates large distance). Typically, one pair of datasets is distinctly more similar than all other pairs.

To demonstrate feasibility and assess the performance of our method, we carried out an in-silico multiplexing experiment on targeted scDNA-seq dataset from 24 acute myeloid leukaemia patients generated from MissionBio’s Tapestri^®^ platform [24], according to an optimal 4-scheme. From this, we obtained pooled samples with ground truth sample identity for each cell. The simulation is described in more detail in Sub-section 3.4. The pools were demultiplexed and each dataset was annotated by ground truth sample identity based on the sample identity of the majority of cells. After running the sample identification algorithm, we compared the established sample identification mapping with the ground truth mapping by computing 1 minus the fraction of mismatches.

The simulations show that, conditioned on successful demultiplexing, the sample identification problem is solved effectively and reliably across cell counts ranging from 1000 to 4000, and doublet rates between 0% and 8%. The performance of the sample matching drops slightly at cells counts of 1000 and a doublet rate of 2%, likely driven by a higher variance in the estimation of the average genotypes of the demultiplexed datasets (see Figure 7**a**). To showcase the robustness of our method, we simulated robust multiplexing and sample matching for pools of 1000 cells at a maximal pool size of 4 and a doublet rate of 5%. We marked demultiplexing experiments as failed, when two datasets from the same pool received the same ground truth sample label. We compared the performances of sample identification in failed and successful demultiplexing experiments. The sample matching performance remains high (see Figure 7**b**). For example, a score of 0.9 means that 2 out of 20 samples were incorrectly matched. A slight drop in performance upon failure of demultiplexing is to be expected, since our algorithm only establishes one-to-one mappings between datasets and samples, which is not possible when the demultiplexing algorithm creates two clusters with the same ground truth label. To further assess quantitatively how the performance of the demultiplexing affects the sample identification, we plotted the sample identification score against the V-Score, a measure to evaluate the performance of clustering [25] (see Figure 7**c**). Our simulations show that robust sample identification works reliably when the V-Score is above 0.75, a performance that was typically reached in simulation studies by various demultiplexing tools [2].

**Fig. 7.**
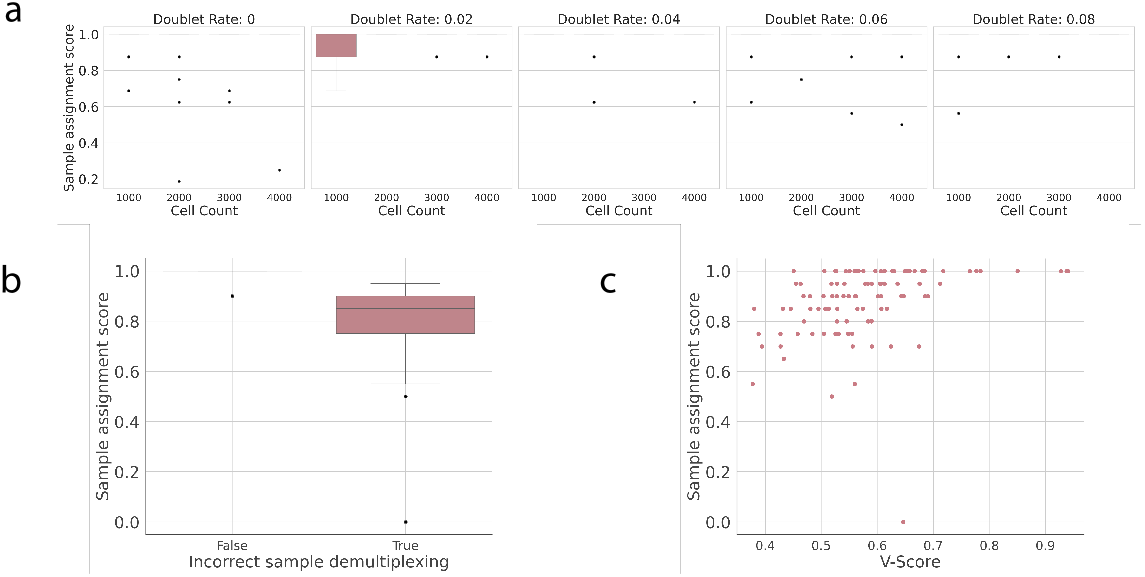
Performance assessment in a simulation study. The performance score is 1 minus the fraction of mismatched samples. A higher score is better. **a**. Boxplots show a high score across different simulated doublet rates and cells counts for all setting for which sample demultiplexing was annotated as successful. **b**. When demultiplexing fails to identify sample clusters correctly, robust pooling strategies maintain sample identification at high performance. **c**. Each point is a simulated run of a robust sample identification. The x-axis shows the V-Score of demultiplexed clusters compared to the ground truth labels. Sample identification becomes more reliable when the demultiplexing performs better.

## 3 Methods

### 3.1 Sample matching

The genetic profiles of demultiplexed datasets are represented as vectors *v ∈* [0, 1]^*N*^, where *N* is the number of loci. Each entry corresponds to the variant allele frequency at a specific position. Two different samples may have information available for different positions. Akin to [2], their genetic similarity is the euclidean distance calculated in the intersection space (on all common positions of the two vectors), divided by the dimensionality of the intersection space.

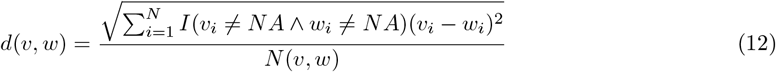

Here, *I* denotes the indicator function, *NA* means that no genotyping information is available for the dataset at a specific genomic position, and *N* (*v, w*) is the dimensionality of the smallest space containing *v* and *w*.

### 3.2 Demultiplexing

For demultiplexing, we use a variant of DemoTape [2]. The distance between two cells is defines as follows. A cell is represented as a matrix *A ∈* N^*N×*2^ where *N* is the number of loci, and for each locus *i, A* contains a pair of non-negative integers *A*_*i*_ = (*a*_*i*_, *t*_*i*_) indicating the alternative allele counts and total allele counts, respectively. Alternative read counts at a specific locus are modelled by a binomial distribution, i.e., *a*_*i*_ *∼ B*(*t*_*i*_ *θ*). We assume independence between sites. In summary, we define the likelihood of the read counts of a cell as

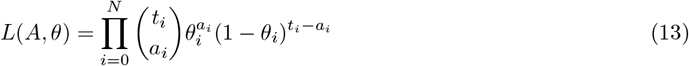

To obtain a genetic distance between two cells, we compare the two models where the alternative counts are sampled from total counts in both cells independently, and where, at each locus, the alternative read counts are sampled from the total read counts with the same probability in both cells. The distance of cells *A* and *B* is proportional to the likelihood ratio

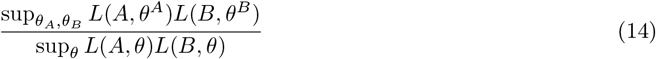

and normalised such that the maximal distance is equal to one.

After calculating the pairwise distances of cells, we use hierarchical agglomerative clustering on all cells with Ward linkage [33]. When *k* samples are expected in the pool, *k* + 1 many clusters are identified, and the smallest cluster is assumed to consist of doublets and removed from the dataset.

### 3.3 Performance assessment

To assess the performance of the sample identification, each demultiplexed cluster was annotated with its sample identity by the majority ground truth label of the majority of cells. Simulated demultiplexing results were flagged as incorrect, if two demultiplexed datasets received the same sample label, so as to not confound the performance assessment of the sample identification by the performance of the demultiplexing step. For each simulation run in each condition, the number of incorrectly labelled datasets was counted and normalised by the total number of datasets as a performance metric.

### 3.4 Simulation study

The patient cohort was filtered to only contain one sample per donor to guarantee that two samples in a pool would never originate from the same donor. For each simulation run, samples were uniformly chosen from the cohort, the cells were randomly split into two datasets with probability sampled from a beta-binomial distribution with shape vector (6, 6). Pooling was performed in-silico using the benchmarking pipeline from [2], simulating unequal proportions by randomly sampling from a Dirichlet-distribution with all shape parameters equal to 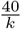, where *k* is the maximal pool size. We downsampled the multiplexed datasets to reach a total cell count ranging from 1000 to 4000 cells and simulated doublet rates ranging from 0% to 8%.

## 4 Discussion

We have presented a novel class of procedures to efficiently multiplex samples and identifying samples in each demultiplexed dataset without the need for additional genotyping or for labelling of samples with hashtag oligonucleotides. Our approach takes into account the constraint that the maximal number of samples to be multiplexed is limited by the capability to computationally demultiplex the sequenced samples. We developed a mathematical formalism to describe this task and constructed an optimal pooling strategy for all sample sizes and all upper bounds on the number of samples per pool. Furthermore, we provide a robust procedure which allows the reconstruction of the sample identity even when the data or parts of the data from one pool become unavailable. This approach takes into account the possibility that complex pooling strategies may lead to accidentally incorrect execution of the laboratory work and the possibility of computational errors during demultiplexing. Our method enables single-cell sequencing to be performed at a substantially reduced cost and facilitates its application in research and the clinics.

One limitation of our approach is the need to correctly identify pairs of samples of identical genotypes. Sequencing data is inherently noisy and the purity of the demultiplexed samples is impacted by the performance of the demultiplexing method. These distortions of the genotype profiles may lead to incorrect or impure demultiplexed samples and downstream to wrong sample labelling. However, genotype demultiplexing methods exhibit correctness and a high purity of clusters in simulation studies, such that for practical purposes, a high reliability of the sample identification is expected. Another practical limitation is due to the fact that library preparation is a labour intensive task and samples typically need to be processed fast. Splitting samples into different pools therefore requires the preparation of several pools simultaneously or within a short amount of time.

Other demultiplexing methods rely on additional genotype profiling of the donors or indexing of reads with hashtag oligonucleotides by sample. These methods typically allow experiments pool by pool. While in some cases those approaches enable a joint library preparation of a higher number of samples per pool compared to the approach proposed here, they suffer from drawbacks: In order to be able to perform additional genotype experiments, more sample material must be available. However, tissue material may be very limited, for example, when samples were collected in a biobank prior to realization of a study. Using hashtag oligonucleotides to label reads by patients requires the design of appropriate primers. For example, hashtag oligonucleotide cocktails compatible with MissionBio Tapestri^®^ reagents have been designed by BioLegend under the name TotalSeq™-D, but only three different hashtag oligos are currently available off the shelf, limiting the pool size to three. By contrast, our combinatorial pooling approach allows sample identification in the absence of any additional genotyping or indexing data.

### Implementation

The code and all simulations have been implemented in python and snakemake [22] and are freely available under a GPL3 license at https://github.com/cbg-ethz/mixmax.

## Data availability

No original data was produced in this study. The data was published by [21] and the raw data is available on the SRA under the project ID PRJNA648656 and the authors shared a subset of the preprocessed loom files with us.

## Author contributions

J.G., N.Bo. and N.B conceptualised the study. J.G. performed the formal analysis, investigation, validation, visualisation, and wrote the first draft of the manuscript. J.G. and N.Bo. provided code. J.G. and L.C. developed the methodology. N.B. and L.C. acquired the funding, N.B. administered and supervised the project. J.G., L.C. and N.B. edited the manuscript. All authors reviewed and approved the manuscript.

## Acknowledgments

We thank Christian Beisel for his insightful comments. L.C. was funded by FAPERJ-JCNE (E-26/201.372/2022) and CNPq-Universal (406173/2021-4). J.G. was funded by the LOOP Zurich (INTeRCePT). Part of this work was funded by the EC Horizon 2020 program (project OLISSIPO, No. 951970).

