## Supplementary Information for "Robust and cost-efficient single-cell sequencing through combinatorial pooling"

### 1 Proofs of Theoretical Results

In this section we will provide proofs for all theoretical results in the main manuscript.

**Theorem 1.** *The highest achievable efficiency for a $k$-scheme is*

$$e(k) = \frac{k+1}{2}. \tag{1}$$

*Furthermore, there is an optimal $k$-scheme with $k$ pools and $\frac{k(k+1)}{2}$ many samples and it can be constructed explicitly.*

*Proof.* First, we show that $\frac{k+1}{2}$ is an upper bound on $e(k)$. Let $\Phi : S \to \mathcal{P}(\Omega)$ be an arbitrary $k$-scheme. We define $T := \sum_{s \in S} |\Phi(s)|$. By the injectivity of $\Phi$, each one-element subset of $\Omega$ can be mapped to at most once, and all other elements must map to subsets of at least two elements. As a direct consequence,

$$T \geq |\Omega| + (|S| - |\Omega|)2 \tag{2}$$

Furthermore, since $T = \sum_{\omega \in \Omega} \Phi_i^{-1}(\omega)$ and $\Phi_i^{-1}(\omega) \leq k$ for each $i$, it follows that

$$T \leq k|\Omega| \tag{3}$$

Putting the equations together and solving for $e(\Phi) = \frac{|S|}{|\Omega|}$, we obtain $e(\Phi) \leq \frac{k+1}{2}$ and since $\Phi$ was chosen arbitrarily, this implies

$$e(k) \leq \frac{k+1}{2} \tag{4}$$

Next, we show that the upper bound can be attained. Define a set of pools $\Omega$ of size $k$ and a set of samples of size $\binom{k}{1} + \binom{k}{2}$, corresponding to $\binom{k}{1}$ many subset of $\Omega$ of size 1 and $\binom{k}{2}$ many subset of $\Omega$ of size 2. Hence, we can choose an arbitrary bijective map between $S$ and the set of nonempty subset of size at most 2 of $\Omega$. By construction this is a multiplexing scheme.

It remains to prove that $\Phi_i^{-1}(\omega) \leq k$ for all $\omega \in \Omega$. By construction, there is exactly one element in $S$ mapping to $\{\omega\}$ and exactly $k-1$ many 2-element subsets of $\Omega$ containing $\omega$. In particular, $|\Phi_i(\omega)^{-1}| = k$. This shows that $\Phi$ is indeed a $k$-scheme.

Finally, note that $e(\Phi) = \frac{|S|}{|\Omega|} = \frac{\binom{k}{1}+\binom{k}{2}}{k} = \frac{k+1}{2}$, which completes the proof. □

**Theorem 2.** *Let* $k \in \mathbb{N}_{\geq 1}$, $n \in \mathbb{N}_{\geq 1}$ *and* $n > \frac{k(k+1)}{2}$.

1. *Let $k$ be odd. Then the smallest number of pools to sequence $n$ samples in a robust $k$-scheme is given by*
$$p_n(k) = k + q + 1 \tag{5}$$
*where $q$ is the quotient to the Euclidean division equation with remainder $r$:*
$$n - \frac{k(k+1)}{2} - 1 = q\frac{k+1}{2} + r \qquad \left(q, r \in \mathbb{N}; \quad 0 \leq r < \frac{k-1}{2}\right)$$
2. *Let $k$ be even. Then the smallest number of pools to sequence $n$ samples is given by*
$$p_n(k) = k + 2q + t \tag{6}$$
*where $q$ is the quotient of the Euclidean division equation with remainder $r$:*
$$n - \frac{k(k+1)}{2} - 1 = q(k+1) + r,$$
*where, if $r < \frac{k-2}{2}$ then $t = 1$, otherwise $t = 2$.*

*Consequently,*
$$\max\{e(\Phi) \mid \Phi : S \to \mathcal{P}(\Omega) \textit{ is a } k\textit{-scheme and } |S| = n\} = \frac{n}{p_n(k)} \tag{7}$$

*Proof.* For the first statement, we proceed by induction over $q$.

q=0: By definition, $\frac{k(k+1)}{2} < n$, so $p_n(k) > k$, because otherwise there would be a $k$-scheme $\Phi$ with $e(\Phi) = \frac{n}{p_n(k)} > \frac{k+1}{2}$ which contradicts Theorem 1. Furthermore, assume $n = \frac{k(k+1)}{2} + \frac{k+1}{2}$. We will construct a $k$-scheme with $k+1$ pools. This will prove that $p_n(k) \leq k+0+1$ for the given $n$, and hence also for all $n < \frac{k(k+1)}{2} + \frac{k+1}{2}$, i.e., for all positive $n$ such that $q = 0$. Consider an optimal $k$-scheme $\Phi : S \to \mathcal{P}(\Omega)$ with $k$ pools and $\frac{k(k+1)}{2}$. Such a scheme exists by Theorem 1. We construct another $k$-scheme
$$\Phi' : S \cup \{s_1, \ldots, s_{\frac{k+1}{2}}\} \to \mathcal{P}(\Omega \cup \{\omega_0\})$$

by adding another pool and $\frac{k+1}{2}$ more samples. We define $\Phi'(s_1) = \{\omega_0\}$. By the injectivity of $\Phi$ and the assumption that no element maps to the empty set, We can pick $\frac{k(k-1)}{2} > k-1$ many samples $s$ such that $|\Phi(s)| > 2$. Without loss of generality, assume $\Phi(s) = \{\omega_1, \omega_2\}$. We can define $\Phi'(s) := \{\omega_1, \omega_0\}$ and $\Phi'(s) := \{\omega_2, \omega_0\}$. Applying this procedure to all $\frac{k-1}{2}$ samples and defining $\Phi'(s) := \Phi(s)$ for all remaining samples results in a $k$-scheme with $k+1$ pools. In particular, this scheme is optimal again. Lastly, this $k$-scheme has again more than $k-1$ many elements mapping to non-singleton sets.

q>0: The inductive step is identical. The main observation to be made is that adding one pool allows to add $\frac{k+1}{2}$ many samples.

This completes the proof of the first statement.

The proof of the second case is similar to the first case. □

**Theorem 3.** *Let $k \in \mathbb{N}_{\geq 1}$. The best achievable efficiency for a robust $k$-scheme is*

$$\max\{e(\Phi) \mid \Phi \textit{ is a robust } k\textit{-scheme}\} = \frac{k}{2} \tag{8}$$

*Proof.* Let $\Phi$ be an arbitrary robust $k$-scheme, and for any $\omega \in \Omega$, let $\hat{\Phi}$ be the $k$-scheme obtained from $\Phi$ by discarding $\omega$. First, we establish an upper bound on the efficiency of $\Phi$. Then, we show that this upper bound is attained by constructing an optimal robust $k$-scheme.

Any robust scheme satisfies $|\Phi(s)| \geq 2$ for all $s \in S$. Assume the contrary. No element maps to the empty set in a multiplexing scheme. So there is some $s_0 \in S$ for which $\Phi(s_0) = \{\omega_0\}$, in which case $\hat{\Phi} : S \to \Omega \backslash \{\omega_0\}$ maps $s_0$ to the empty set, contradicting the definition of a robust $k$-scheme. All elements $s \in S$ mapping to a singleton set through $\hat{\Phi}$ satisfy $\Phi(s) = \hat{\Phi}(s) \cup \{\omega\}$, and since $\Phi$ is a $k$-scheme we have $\Phi^{-1}(\omega) \leq k$. Hence, there are at most $k$ many elements mapping to a singleton through $\hat{\Phi}$. Akin to the proof of Theorem 1 we define $\hat{T} := \sum_{s \in S} |\hat{\Phi}(s)|$ and see that

$$k + 2(|S| - k) \leq \hat{T} \leq (|\Omega| - 1)k \tag{9}$$

which, solving for $e(\Phi) = \frac{|S|}{|\Omega|}$, yields

$$e(\Phi) \leq \frac{k}{2} \tag{10}$$

Next, take an optimal $k$-scheme $\hat{\Phi} : S \to \mathcal{P}(\hat{\Omega})$ such that $|\hat{\Omega}| \leq k$ as constructed in Theorem 1. We construct a robust $k$-scheme by adding an additional pool $\omega$ as follows:

$$\Phi : S \to \mathcal{P}(\hat{\Omega} \cup \{\omega_0\}), \quad s \mapsto \begin{cases} \hat{\Phi}(s) & \text{if } |\Phi(s)| = 2 \\ \hat{\Phi}(s) \cup \{\omega_0\} & \text{otherwise} \end{cases} \tag{11}$$

$\Phi$ is injective because $\hat{\Phi}$ is, and no element maps to the empty set. Furthermore, at most $|\Omega|$ many samples map to singleton sets, so $|\Phi^{-1}(\omega_0)| \leq |\Omega| \leq k$, and for any other $\omega \in \Omega$, we have $\Phi^{-1}(\omega) = \hat{\Phi}^{-1}(\omega)$ which has at most $k$ elements. This proves that $\Phi$ is a $k$-scheme. To show that it is robust, first note that discarding $\omega_0$ yields the $k$-scheme $\hat{\Phi}$ , so we only need to show that $\hat{\Phi}' : S \to \Omega \backslash \{\omega\}$ is a $k$-scheme for $\omega \in \hat{\Omega}$ . Since $|\Phi(s)| \geq 2$ for all $s \in S$ by construction, $\hat{\Phi}'$ maps no element to the empty set.

Next, we show that $\hat{\Phi}'$ is injective. So let $s_1, s_2 \in S$ and $s_1 \neq s_2$, and assume that $\hat{\Phi}'(s_1) = \hat{\Phi}'(s_2)$. We will lead this to a contradiction. If $\omega_0 \in \hat{\Phi}'(s_i)$, $(i = 1, 2)$, then $\hat{\Phi}'(s_i) = \{\omega_0\}$ or $\hat{\Phi}'(s_i) = \{\tilde{\omega}, \omega_0\}$ for some $\tilde{\omega}$, so $\hat{\Phi}(s_1) = \{\omega, \omega_0\} = \hat{\Phi}(s_2)$ or $\hat{\Phi}(s_1) = \{\tilde{\omega}, \omega_0\} = \hat{\Phi}(s_2)$, and both cases imply $s_1 = s_2$. If $\omega_0 \notin \hat{\Phi}'(s_i)$, then $\hat{\Phi}'(s_i) = \Phi(s_i)\{\omega\}$ and both $\Phi(s_i)$ map to 2-element subsets of $\Omega$, so $\Phi(s_1) = \Phi(s_2)$ and therefore $s_1 = s_2$. This proves that $\hat{\Phi}'$ is injective. Thus, $\Phi$ is a robust $k$-scheme.

Since $\hat{\Phi}$ is optimal, its efficiency is equal to $\frac{k+1}{2}$ (by Theorem 1), and therefore $|S| = \frac{(k+1)|\hat{\Omega}|}{2}$. As a consequence,

$$e(\Phi) = \frac{(k+1)|\hat{\Omega}|}{2(|\hat{\Omega}|+1)} = \frac{k+1}{2} - \frac{k+1}{2(|\hat{\Omega}|+1)} \leq \frac{k}{2} \tag{12}$$

since we have assumed $\hat{\Omega} \leq k$. □
